# Performance Assessment of an Ultraviolet Light Emitting Semi-Conductor Device in Treating Apple Juice: Microbial Inactivation and Biochemical Assessment Study

**DOI:** 10.1101/2022.10.11.511833

**Authors:** Anita Scales Akwu, Ankit Patras, Brahmiah Pendyala, Anjali Kurup, Fur-Chi Chen, Matthew J. Vergne

**Affiliations:** Department of Agricultural and Environmental Sciences, Tennessee State University, Nashville, TN 37209, USA; Department of Pharmaceutical Sciences and Department of Chemistry & Biochemistry, Lipscomb University, Nashville, TN 37204, USA

## Abstract

Inactivation of *Listeria monocytogenes* ATCC 19115 and *Salmonella enterica* serovar Muenchen ATCC BAA 1764 by a light emitting diodes (LED) operating at 279 nm was investigated. In addition, this investigation assessed the poly-phenolic and vitamin content of UV irradiated apple juice (AJ). Specific concentrations of bacteria were inoculated in AJ and irradiated at the designated UV doses of 0 to 10 mJ·cm^-2^ for *Salmonella* Muenchen and 0 to 12 mJ·cm^-2^ for *Listeria monocytogenes*.Results show that UV-C irradiation effectively inactivated pathogenic microbes in AJ. The log reduction kinetics of microorganisms followed log-linear and with higher R^2^ (>0.95). The D_10_ values of 3.50 and 3.56 mJ·cm^-2^ were obtained from the inactivation of *Salmonella* Muenchen, and *Listeria monocytogenes* in apple juice. In addition, quantifiable UV-C doses ranging from 0 to 160 mJ·cm^-2^ were also delivered to AJ and polyphenols and vitamins were profiled. LC-MS/MS analysis was conducted to assess the stability of polyphenols or vitamins in UV-C exposed AJ. The polyphenol and vitamin results demonstrated that UV-C irradiation in AJ can cause significant reductions (p<0.05) if not properly delivered. Chlorogenic acid was reduced to 56%, at 80 mJ/cm^2^ whereas 12% reduction was observed at 40 mJ/cm^2^. Choline was observed to be relatively stable as a function of UV-C dosage. In contrast thiamine was significantly reduced at higher doses. In addition, Epicatechin was significantly reduced at high exposure doses. In contrast minor changes were observed at 40 mJ/cm^2^. The results from this study imply that adequate log reduction of pathogens is achievable in AJ and suggest significant potential of using LED devices for UV-C treatment of highly turbid fluids.

## 1. Introduction

Apples and their respective products contain numerous antioxidant phytochemicals which exhibit bioactive properties. Phytochemicals are best defined as nutrient bioactive chemicals derived from plants present in fruits, vegetables, and grains; and provide advantageous benefits to overall health outside of basic nutrition to reduce major chronic disease risks (Jimenez-Garcia, et al., 2018; Liu, 2004). The bioactive compound categories include phytochemicals, vitamins, minerals, and dietary fibers. Apple juice is also composed of a variety of sugars, which primarily include, fructose, glucose, and sucrose (Pina-Pérez, Rodrigo & Martinez, 2015). In addition to these sugars, is also the presence of starches (oligosaccharides and polysaccharides), acids (malic, quinic, citramalic, organic, amino), tannins (polyphenols, phenols), amides and other nitrogenous compounds, soluble pectin, vitamin C, minerals, boron, and a vast array of esters (Pina-Pérez, Rodrigo & Martinez, 2015; Smeriglio, et al., 2017; Swamy, Muthukumarappan, & Asokapandian, 2018, Wojdyło, et al., 2021; Teleszko & Wojdyło, 2015).

Apple juice is heavily consumed because of the many nutritional benefits (Muñoz, et al., 2012), pleasant organoleptic qualities (Muñoz,et al., 2012; Włodarska, et al., 2019), prebiotics content (Ribeiro et al., 2021), probiotics (Dimitrovski, et al., 2015; Cousin, et al., 2017), polyphenols (Du, et al., 2019; Zhang, et al., 2021), vitamins A,C, E, K (Islam, et al., 2016b; Karasawa & Mohan, 2018; Akwu, et al., 2022), minerals and dietary fibers (Karasawa & Mohan, 2018), strong antioxidant activity, cancer cell proliferation inhibition, lipid oxidation decreases, and lowered cholesterol (Akwu, et al., 2022; Kidoń & Grabowska, 2021; Boyer & Liu, 2004) contents. Collectively, they function in providing great benefits to our bodies against diseases, ailments, clinical anti-allergic activity (Heinmaa, et al., 2016). Perhaps due to high sugar content and high vitamin levels, AJ can be contaminated with vegetative cells, spores and other micro-organisms.

Fruit juices can be spoiled due to the growth of microorganisms. Yeasts and molds, *Lactobacillus, Leuconostoc* and thermophilic *Bacillus* are common spoilage microorganisms. Perhaps the juices can contain pathogenic micro-organism at 1-2 log_10_ concentration level. Control measures for low acid and acidic beverages are critical, and are likely to involve multiple measures, for example, a combination of a process steps to destroy the nonproteolytic spores and ‘‘Keep Refrigerated” labeling if the juice does not receive a treatment sufficient to destroy the proteolytic spores (21 CFR Parts 113 and 114). Additionally, guidance from the FDA is now strictly recommending that processors subject to the pathogen reduction provisions of the juice Hazard Analysis and Critical Control Points (HACCP) regulation (USFDA, 2004) incorporate validated control measures for all *B. cereus* and *C. botulinum* spores into their HACCP plans (USFDA, 2007), to control their growth and toxin production. For acidic juice products, *Escherichia coli, Saccharomyces cerevisiae and Listeria innocua* are associated with food-borne pathogens (Basaran, Quintero-Ramos, Moake, Churey, & Worobo, 2004; Guerrero-Beltrán & Barbosa-Cánovas, 2005). A 1998 FDA ruling Code of Federal Register, (63 FR 37030), made it mandatory for fruit and juice processors to provide potential sickness product labels if consumed without a 5log_10_ reduction of pathogenic microflora. As a result of this ruling, processors began utilizing conventional heat methods such as pasteurization to reduce the microbial population by a 5-log_10_ reduction. In addition to this rule, the U.S. Food and Drug Administration suggested that retailers also implement the 5-log_10_ reduction in addition to their respective state laws and regulations to be in compliant. Heat severely impacted the quality of juice products. Several different authors reported the impact of heat on vitamins and polyphenols (Patras, et al., 2020; Patras, Tiwari, & Brunton, 2011; Akwu, et al., 2022). As a result of the heat impact on product quality, it is important to explore alternative methods of pasteurization and sterilization that will retain the quality and safety of the product.

A document titled ‘Kinetics of Microbial Inactivation for Alternative Food Processing Technologies: Executive Summary; was published by Food and Drug Administration and the Center for Food Safety and Applied Nutrition. This report evaluates and elaborates on alternative technologies and addresses the knowledge gaps and research needs. (CFSAN, 2000). According to *Akwu, et al. 2022*; there is an urgent to investigate novel non-thermal processing technologies and generate microbial inactivation for a range of pathogens to demonstrate its effectives. UVLEDs are high-tech technologies that require validation via scientific research studies to demonstrate, “proof of principle;” which demonstrates the efficacy of them as a novel technology (Akwu, et al., 2022). UV technologies have shown considerable promise for pasteurization and sterilization of acidic and low acid fluids. (Akwu, et al., 2022; Patras, et al., 2020; Delorme, et al., 2020; Rahman, 2020; Koutchma, 2019; Islam, et al., 2016b). These alternative pasteurization methods have been registered for juices through the Code of Federal Register (CFR 179.39).

The electromagnetic spectrum contains three different wavelengths for UV-C irradiation that range from 100 nm to 400 nm. Between 320 nm and 400 nm is classified as the UV-A range, UVB is classified between 280 nm and 320 nm, and the UV-C range is classified between 200 nm and 280 nm (Akwu, et al., 2022; Centers for Disease Control and Prevention, 2022; Johns Hopkins Medicine, 2022; World Health Organization, 2016). Each of the wavelengths A, B, and C can pose challenges to our overall health, if we do effectively prepare ourselves. UV-C is the most effective antiseptic or disinfectant agent against many different microorganism types; as photons are consumed by DNA (Akwu, et al., 2022; Yang et al., 2019; Yin et al., 2013; Dai et al., 2012). As Patras, et al., 2020 mentioned, photochemical reactions that take place in DNA function by hindering the ability of microbial replication to occur. UV-C irradiation studies uses low pressure lamps as an optical source. There is an interest in implementing LED device for disinfection studies (Akwu, et al., 2022).

LED devices utilizing high UV intensity have been utilized by several authors (Kurup, et al., 2022; Akwu, et al., 2022; Prasad, et al., 2020; Kusuma, Pattison, & Bugbee, 2020). Though many studies have utilized ultraviolet (UV) light emitting diodes (LEDs); two major weaknesses are present to include, verification and calculation of the UV dosage (Heckman, et al., 2013; Beck et al., 2017) and the inadequacy of optical data utilized in the studies (Akwu, et al., 2022; Caminiti, et al., 2012; Unluturk, et al., 2010). LED spectral output is not a single wavelength but a collection of other wavelength (±10 nm), average fluence calculations need to account for all wavelengths. This study fills knowledge gaps in the literature on the efficacy of LED’s and its ability to inactivate pathogens at relevant inactivation doses.

The main objective of this study was to develop standardized UV fluence response curves for *Listeria monocytogenes* ATCC 19115 and *Salmonella enterica* serovar Muenchen ATCC BAA 1764 suspended in apple juice under dynamic stirred conditions. A secondary aim was to assess the effect of UV-C irradiation on polyphenols and vitamins in apple juice.

## 2. Materials and methods

### 2.1 Chemicals

Phloridzin dihydrate, was bought from Sigma Aldrich, USA, Whereas, (-)-epicatechin was bought from Adooq Bioscience, CA, USA. Furthermore, analytical grade epicatechin, thiamine, choline chloride, B12 and Phlorizin were purchased from Sigma Aldrich, Missouri, USA. HPLC and LCMS grade water, methanol and acetonitrile were sourced from Fisher Scientific, USA.

### 2.2 Sample Preparation

Similar preparation was conducted as described by Akwu et al., 2022. Apples (*cv* Gala) were obtained from a local grocery store. All apples were thoroughly washed and juiced using a Brentwood 800W juicer. The juice was strained and then filtered using Whatman (28-30 μm, 2-3 μm). Optical data was obtained and recorded and 45mL per culture tube were stored at −20°C wrapped with aluminum foil until further processing, to avoid exposure to light. Two filtration steps were implemented to remove solid particulates. Prior to irradiation, AJ samples were randomly assigned to a treatment process and thawed at room temperature. Following the juicing process, baseline optical data was obtained, and apple were divided into 20 aliquots of 10 ml each stored at −20 °C until used. Physicochemical characteristics of the AJ is shown in Table 1. A balanced design with three replicates randomized in experimental order was performed for each UV dose.

**Table 1.** Physiochemical characteristics of apple juice.

| **Parameters** | **Values** |
| --- | --- |
| pH | $3.3 \pm 0.164$ |
| Total soluble solids (%) | $14.3 \pm 0.018$ |
| Titrateable acidity | $5 \pm 0.406$ |

Data presented as mean ± stdev

### 2.3 Bacterial strains and cultural conditions

Two non-pathogenic and non-outbreak strains of different bacteria were used in this study *Salmonella enterica* serovar Muenchen ATCC BAA (1764) and *Listeria monocytogenes* (19115). The bacterial strains were procured from American Type Culture Collection (ATCC). The bacterial cultures were stored in 25% glycerol in cryovials at −80 °C. Fresh bacterial suspensions were prepared for inoculation into apple juice for every treatment. Two loops of individual strains of *S.* Muenchen *and L. monocytogenes* were transferred to 15 mL Tryptic soy broth (Oxoid Ltd., Basingstoke, UK) and incubated at 37 °C for 18 h. *L. monocytogenes* was also subjected to two successive transfers in tubes containing 15 mL of Buffered Listeria enrichment broth (Oxoid Ltd., Basingstoke, UK) and incubation was done for 24 h at 37 °C. These cultures were used as the adapted inoculum. After incubation, *S*. Muenchen culture was transferred into 15 mL of TSB and incubated for 18 h at 37 °C to reach the stationary growth phase. Similarly, *L. monocytogenes* culture was transferred to 15 mL *Listeria* enrichment broth (Oxoid Ltd., Basingstoke, UK) and incubated for 24 h at 37 °C. Centrifugation (3000 × g, 15 min) was done to harvest the bacterial cells. A solution of 0.1% (w/v) phosphate-buffered saline (PBS), Becton Dickinson, New Jersey, US) was used to wash the cell pellets and re-suspended in 50 mL of PBS. For determining the original cell population densities, appropriate dilutions of each cell suspension were made in 0.1% peptone water (PW) and plated in duplicate using Tryptic Soy Agar (Oxoid Ltd., Basingstoke, UK) plates for *S*. Muenchen suspensions and incubation was done at 37 °C for 24 h. *L. monocytogenes* suspensions were plated on Listeria selective agar base (SR0141E) (Oxoid Ltd., Basingstoke, UK) plates with incubation at 37 °C for 48 h.

### 2.4 Apple juice inoculation

Aliquots of 45 mL of AJ were inoculated individually with each of the two bacterial cultures (*L. monocytogenes, S*. Muenchen) targeting a concentration of 10^7^ CFU/mL. The inoculated apple juice was plated using decimal dilutions on Tryptic soy agar (Oxoid Ltd., Basingstoke, UK) plates to determine the original *S*. Muenchen titers and incubation was done at 37 °C for 24 h. Apple juice inoculated with *L. monocytogenes* was plated on *Listeria* selective agar base (Oxoid Ltd., Basingstoke, UK) and plates were then incubated at 37 °C for 48 h.

### 2.5 Optical properties and pH measurements

A method described by Akwu et al (2022) was adapted for optical property measurements. The absorption coefficient at 279 nm was determined based on transmittance measurements from a Cary 300 spectrophotometer with a six-inch integrating sphere (Agilent Technologies, CA, US). Baseline corrections, i.e., by zeroing (setting the full-scale reading of) the instrument using the blank and then blocking the beam with a black rectangular slide was carried out. All pH readings were measured using a standard pH meter (Jenway, Staffordshire, UK). All readings were taken in triplicate to lessen the measurement error. From this data, ultraviolet transmittance value was mathematically quantified.

### 2.6 UV-C irradiation treatments

A near collimated beam system that utilizes a light emitting diode device (Irtronix, Torrence, CA, USA) producing irradiation at 279 nm was used for the UV exposures. The system consisted of 3×3 configuration. Electro optical characteristics of UV-C LED has a maximum forward voltage of 35 Vdc, power consumption of 28 Watts with radiant power of 290 mW. To increase mixing, a 5-mL sample was stirred in a10-mL beakers (height: 3.2 cm; diameter: 1.83 cm) (Bolton & Linden, 2003; Chandra, et al., 2017). All optical parameters and UV dose calculations are described in our published studies (Stanley, et al., 2020; Islam, et al., 2016a; Islam, et al., 2016b). Using a high sensitivity sensor (QE Pro series, Ocean Optics, Dunedin, FL, USA), the central irradiance of the UV-C LED system on the surface of the test solution was measured. Based on the central irradiance of the lamp and optical properties (absorption coefficient) of apple juice, the average fluence was calculated. Four correction factors i.e. reflection (RF) factor, petri factor (PF), divergence factor (DF) and water factor (WF) were accounted for in the fluence calculations. UV-C dose was calculated as a product of average fluence and exposure time, equation 2. Correction factors and parameters for obtaining the average fluence rate are shown in Table 2.The average UV fluence rate in the stirred sample can be calculated as (Eqn. 1):

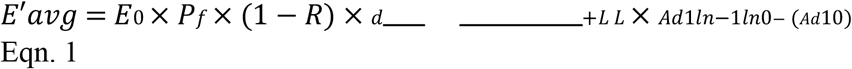

E_0_ is the radiometer meter reading at the center of the beaker and at a vertical position so that the calibration plane of the detector.

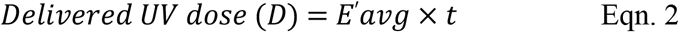

**Table 2.**
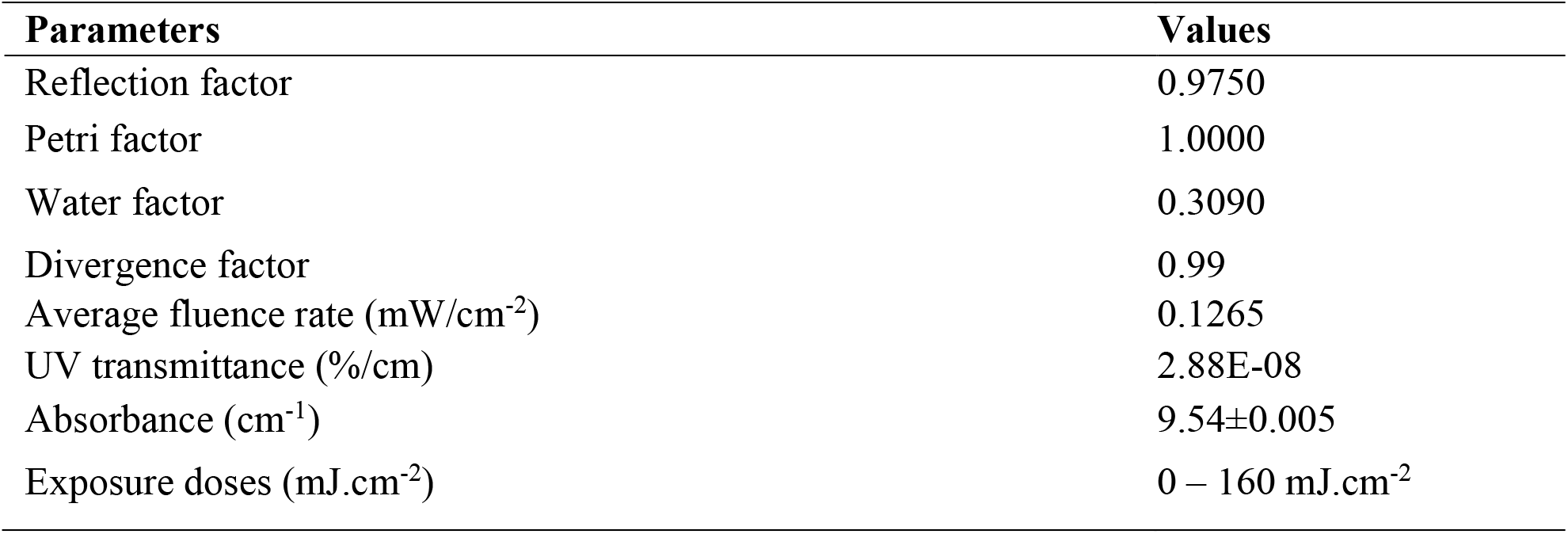
Correction factors and parameters for obtaining the average fluence rate.

### 2.7 LC-MS Methods for Polyphenol Detection

This method was adapted from our previous studies (Akwu et al., 2022). A Shimadzu LCMS 8040 system (Shimadzu Scientific Instruments, Columbia, MD) which included two Shimadzu LC-20ADXR pumps, a SIL-20ACXR autosampler, a CTO-20A column oven, and an LCMS-8040 triple stage quadrupole mass spectrometer was used for LC-MS/MS analysis. Chromatographic separation was achieved on with a Phenomenex 1.6 μm Polar C18 100 Å column (100 × 2.1 mm) maintained at temperature of 40°C. The mobile phase consisted of 0.1% formic acid (FA) in water (A) and 0.1% FA in acetonitrile (B). The flow rate was 0.30 mL/min. Initially, solvent B concentration was 3% and increased linearly to 97% from 0.01 min to 2.0 min. Solvent B was held at 97% from 2.0 to 7.0 min, and then reduced to 3% until the end of the time program at 10.0 minutes. The injection volume was 2 μL. The LCMS analysis utilized an electrospray ionization source with the optimized source parameters DL temperature 250 °C, nebulizing gas flow, 3 L/min, heat block 450°C, drying gas flow, 20 L/min. The following transitions and collision energies (CE) were used to analyze compounds in the positive ion mode: pyridoxal m/z 168>150, CE −13; pyridoxamine m/z 168>152, CE −14; pyridoxine m/z 170>152, CE −15; thioamide m/z 264.9>122.05, CE −15; choline m/z 104.1>60, CE-22, Vitamin B12 m/z 678.2>147, CE −40, and chlorogenic acid m/z 354.9>163, CE −15. These compounds were analyzed in the negative ion mode, ascorbic acid m/z 175>114.85, CE 13; phlorizin m/z 535>273.2, CE 15; and epicatechin m/z 289>125, CE 20. A standard solution of 1000 ng/mL each compound was prepared calibration curves. A one-point calibration curve was used to determine concentrations of samples.

Data were acquired and analyzed with Shimadzu LabSolutions software.

### 2.8 Statistics

The concentration of polyphenols and vitamins were evaluated for six UV doses levels. Microbial studies were conducted for 4 exposure levels. A balanced experimental design with three replicates were randomly assigned to each treatment, i.e. exposure to the selected UV-C irradiation. One-way ANOVA test (Tukey’s HSD multiple comparison) was chosen to analyze the data to evaluate the effects of different UV doses on the concentration of polyphenols and vitamins using SAS computing environment. Data were reported as means ± one standard deviation from the mean and tests were statistically significant at 5% significance level.

## Results and Discussion

From the optical attenuation data (Table 2), it may be seen that the AJ was a strong absorber (7.98 cm^-1^) of UV-C light (Akwu, et al., 2022). Absorbance is a measurement of the amount of UV light that is absorbed by a substance at 279 nm over 1 cm path-length. Fluids exhibiting high absorbance values will create high UV gradients. On the contrary, some fluids have absorbance between value 0.1 to 22 (Patras, et al., 2020; Pendyala, et al., 2021; Vashisht, et al., 2022). Figures 2 and 3 illustrate the ultraviolet (UV) Spectra of apple juice as a function of wavelength. 279 nm wavelengths as demonstrated in Akwu, et al., (2022). Similarly, to, Akwu, et al., (2022), it clearly shows the absorbance at 279 nm is relatively lower in comparison to 254 nm wavelength that demonstrated a higher absorption, which is why 279 nm was the chosen wavelength (Akwu, et al., 2022). UVC exposure of beverages requires very specific requirements to be effective and efficient. One example of UV-C light treatment is a near collimated beam unit that assists in accurate delivery doses of UV-C light in water, wastewater (Gehr, 2007; Kuo, et al., 2003; Qualls, Flynn & Johnson, 1983, Patras et al., 2022). Similar principle can be applied in beverages. UV-C exposure has been well documented as it is notably changed in the presence of optical attenuation coefficient of the apple juice test fluid (Akwu, et al., 2022). UV-C doses must be uniformly delivered in the test fluid to achieve target log reductions of bacteria (Akwu, et al., 2022; Islam, et al., 2016 b; Patras, et al., 2020). This effectively illustrates that log reduction data demonstrated at various optical properties may only be examined in comparative studies if optical properties and UV dose delivered were both accurately accounted for (Patras et al., 2020) As described in Akwu, et al., 2022, all UV gradients in the system were considered in the UV dose/fluence calculations (Akwu, et al., 2022).

**Figure 1.**
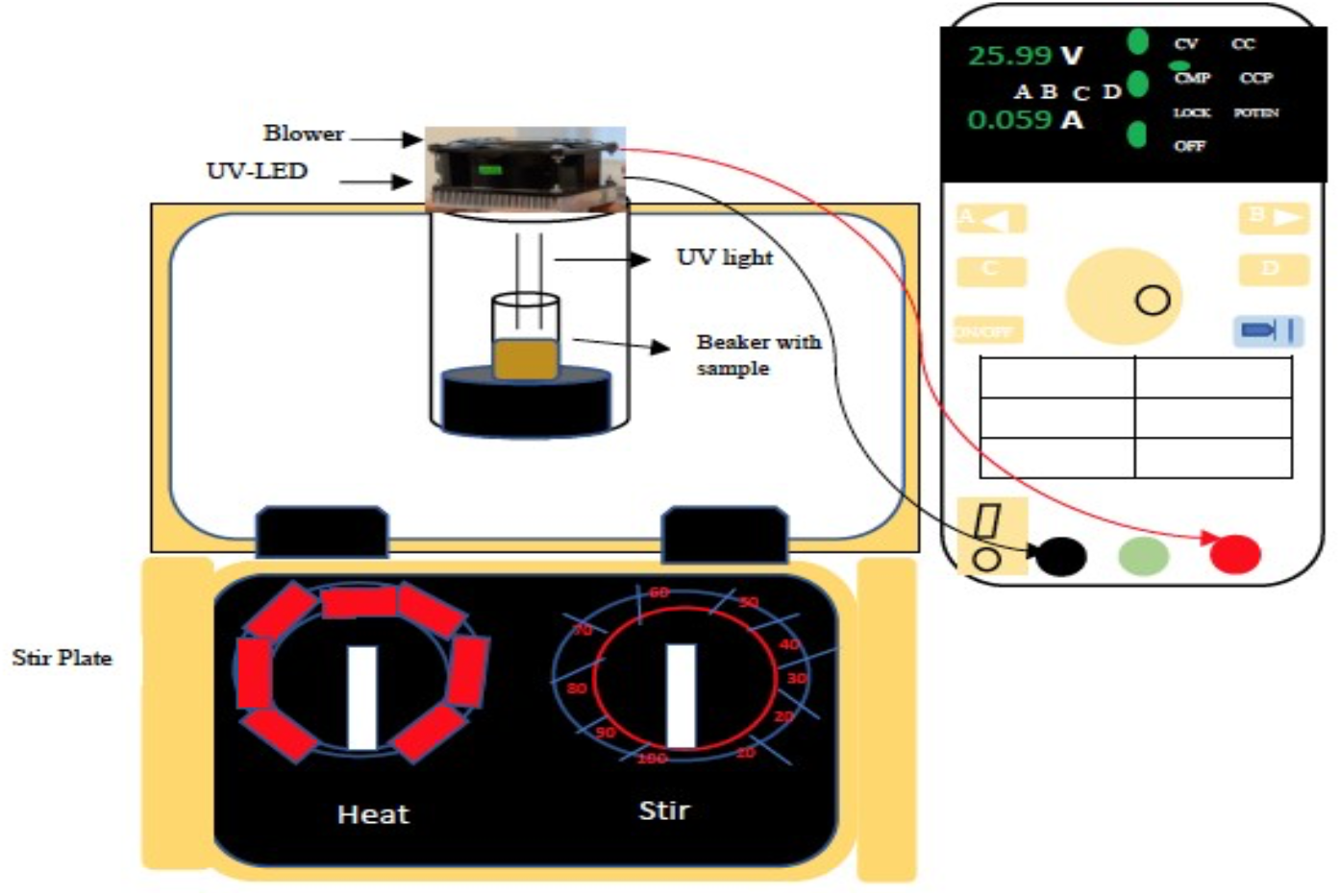
Collimated Light Emitting Diode UV system operating at 279 nm wave-length

**Figure 2.**
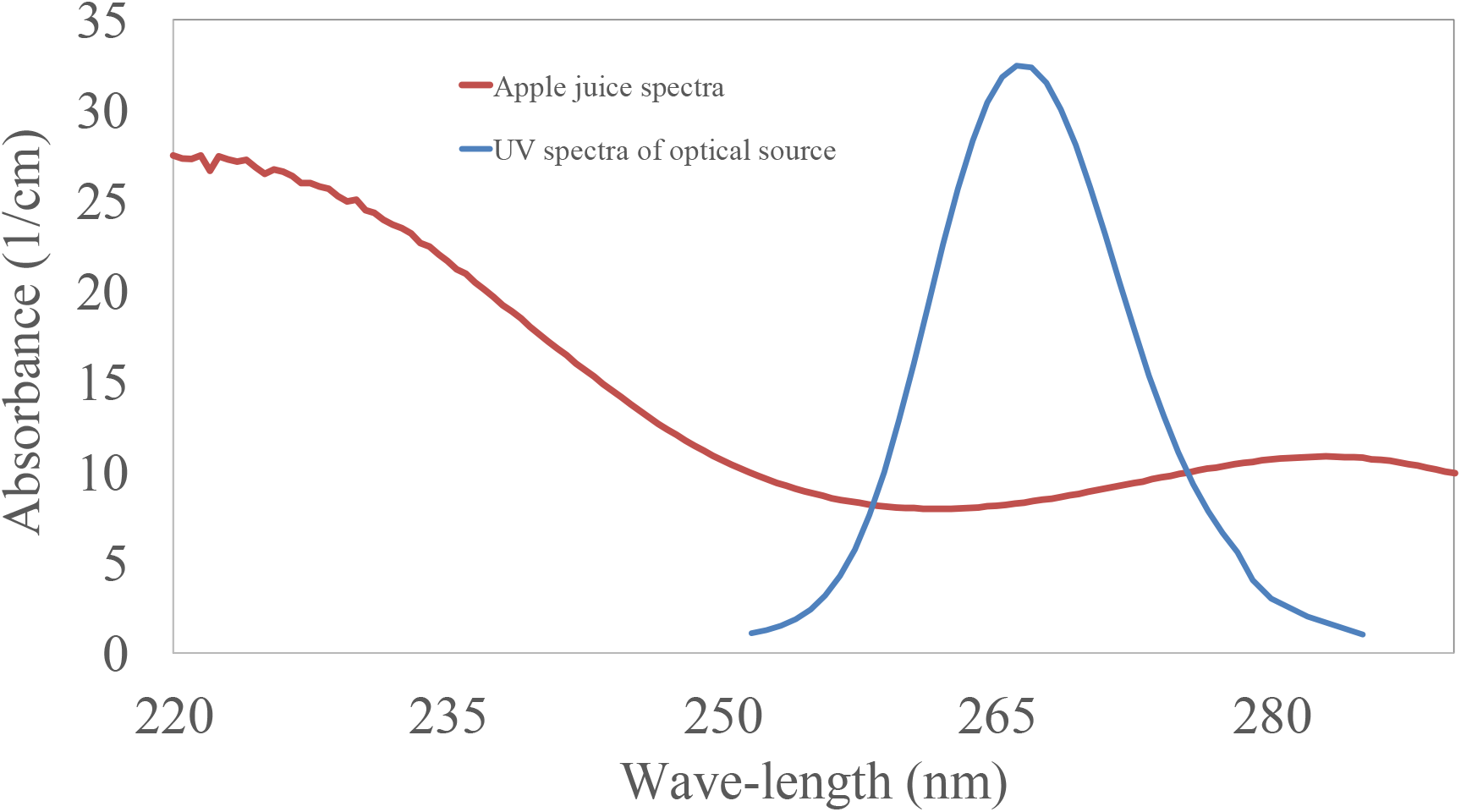
UV Absorbance full scan spectra of apple Juice between 265nm. UV intensity spectra of Light Emitting Diode UV system, triplicate UV scans were performed all replicates shown on plot

**Figure 3:**
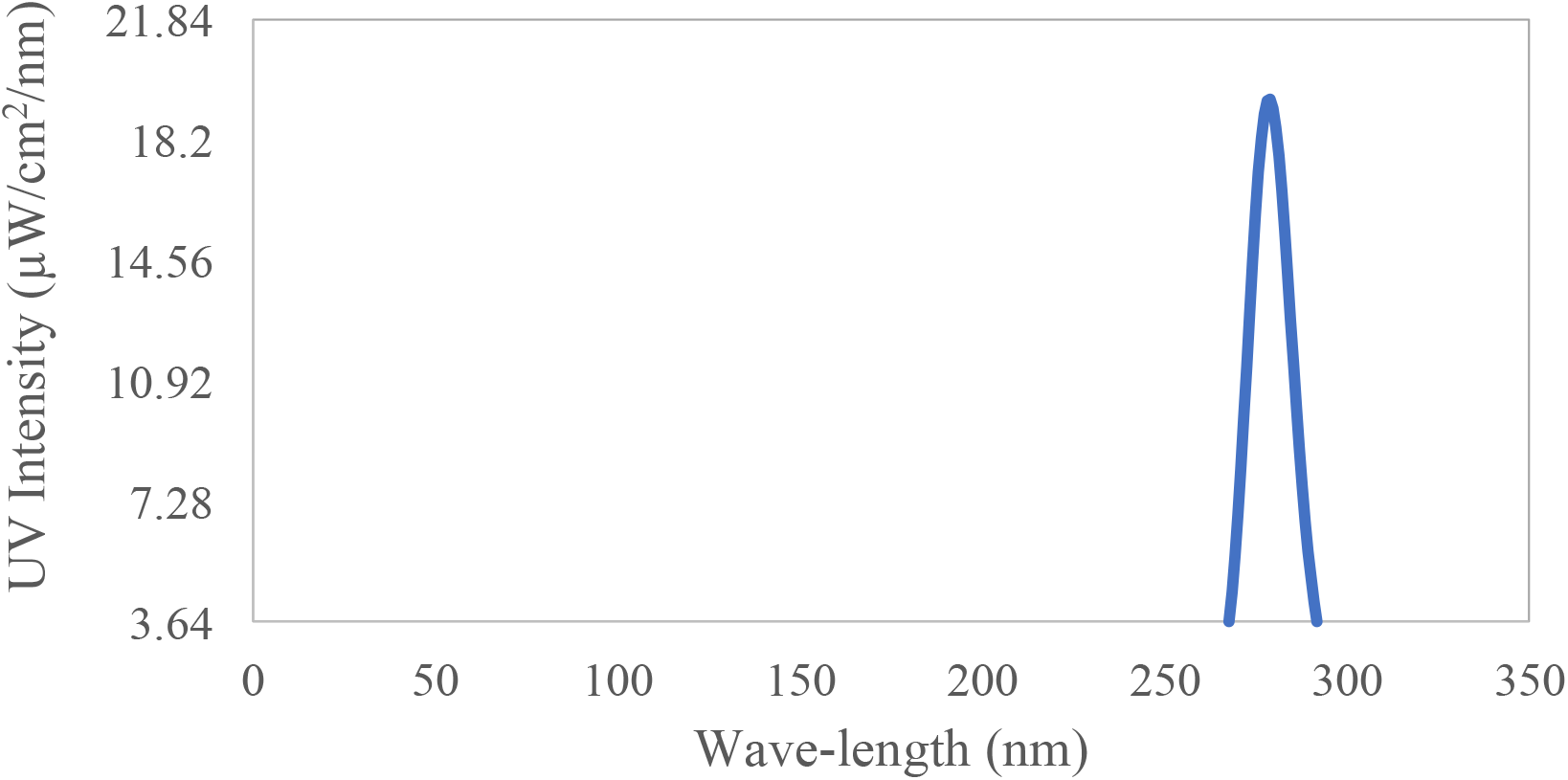
UV Absorbance Scan concentrated around 279nm. UV intensity spectra of Light Emitting Diode UV system, triplicate UV scans were performed one scan shown on plot

UV-C inactivation of different pathogens has been extensively conducted by many researchers (Tosa & Hirata, 1999; Yaun, et al., 2003; Sommer, et al., 2000; Wilson, 1992; Wu, et al., 2011). A great example was demonstrated in McSharry et al, (2022), where *Listeria monocytogenes* T1093, illustrated an initial reduction, then, subsequently first-order kinetics up to 6-log_10_, with D_10_ in the log-linear region of about 1.8 mJ·cm^-2^ (McSharry et al., 2022). Presently, many microbial challenge studies in liquid foods and beverages have been carried out at 254nm (Vashisht, et al., 2022; Sauceda-Gálvez, et al., 2021; Bhullar, et al., 2019; Usaga, et al., 2017; Chandra, et al., 2017; Gunter-Ward, et al., 2017; Torkamani & Niakousari, 2011), however, there is very limited data that is available at the 279 nm wavelength in UV-C microbial challenge studies. We hypothesize that D_10_ value at 254 and 279nm will be very comparable but the optical properties of the suspensions will be significantly different. Kim, Kim, & Kang (2015), studied the effects of Ultraviolet light-emitting diodes (UV-LEDs) on the inactivation of Gram-positive and Gramnegative foodborne pathogenic bacteria and yeasts (Kim, Kim, & Kang, 2015). The bacterial and yeast cocktails were exposed to UVC-LED modules that were connected to electronic printed circuit boards that were clouded for four UV-LEDs with peak voltages at 266, 270, 275, and 279 nm (Kim, Kim, & Kang, 2015). The average UVC-LED voltages applied ranged from 6.36 V to 6.92 V; while the nominal power consumptions were 0.16 W or 0.13 W. The authors observed a significantly higher Propidium Iodide (PI) uptake percentage in Gram-positive in comparison to Gram-negative and Yeasts (Y); which had a maximum of 8% Propidium Iodide uptake (Kim, Kim, & Kang, 2015). It seemed that the authors did not quantify and validate the delivered dose. The authors did not provide the data on fluid mixing.

In a different study, Nyhan et al., (2021), demonstrated that when UVA-LEDS and UVCLEDS could be paired by using the germicidal effect of UV-C and greater penetrating ability of UV-A. The researchers discovered that use of multiple wavelengths 254/270/365 nm demonstrated a reduction after each reduction at 5, 10, 20s, and 40s respectively (Nyhan, et al., 2021). This study investigated the use of UV-LEDs in bacterial inactivation in powdered food ingredients. Damage that occurs to the bacterial membranes is that when exposed to UV-A and UV-C wavelengths microbial inactivation was increased (Chevremont, et al., 2012). Xiang, et al., (2020), applied dosages between 200 - 1200 mJ/cm^2^ and the populations of *Zygosaccharomyces rouxii* in apple juice samples were reduced by 4.86 and 5.46-log at 800 and 1200mJ/cm^2^.

The authors irradiated the samples using a collimated beam apparatus consisted of a low-pressure mercury UV lamp with peak radiation in the 275 nm wavelength range as demonstrated in Fig.1. In our study, 10 mJ/cm^2^ UV dose at 0.126537 mJ/cm2 average irradiance resulted in 3.50 log_10_ CFU/mL reduction of *S*. Muenchen at a depth of 1.5 cm using UV-LEDs emitting light at 279 nm. Fluid absorbance is a critical component in UV technology, as it determines the intensity of UV light and its ability to penetrate. This study rectifies this issue and accounts for UV intensity gradients in the dose calculations (Akwu et al., 2022). The current study examines the inactivation of *Salmonella* Muenchen and *Listeria monocytogenes* in natural apple juice, for which the authors found limited published literature regarding 279 nm wavelength or closer exposure wavelengths.

The populations of *Salmonella* Muenchen were reduced by 1.17, 1.32, 2.90, log_10_ respectively at a UV-C dose level of 4, 5, 10 mJ·cm^-2^. The microbial inactivation results demonstrate first order kinetics, as plotted in Fig. 4. Log inactivation was proportional to UV dosage. Based on the rate constant (cm^2^/mJ) and D_10_ value of *Salmonella*, 17.5 mJ.cm^-2^ dosage will be required to achieve 5 log_10_ reduction. Our experiments were conducted at bench scale and apple juice samples were stirred continuously throughout the irradiation treatment duration, to ensure uniform dose delivery to the homogeneous test fluid (Akwu et al., 2022). The log inactivation increased with increase in the UV-C exposure, as also evidenced in Fig 4. This can be concluded due to the fact that the presence of UV-C light functions in disrupting microbial population replication of DNA and altering gene functions. In some instances when a dose delivery is not uniform, the UV-C dose may exceed the threshold capacity and cellular damage result in rapid lethal inactivation of cells, and DNA repair mechanisms will fail to undo changes (Miller, et al., 1999; Akwu et al., 2022). Previous studies reported D values between 3.5 – 3.85 mJ/cm^2^ for 3 and 4 log_10_ reductions (Gopisetty, et al., 2019; Sommer, et al., 2000); and <2 mJ/cm^2^ through 29 mJ/cm^2^ respectively representing between 1 and 6 total log reduction (Yaun, et al., 2003). All data is in accordance with the published studies except with incidents of high d values, which could be a result of poor mixing within the system. This poor mixing also creates false tailing effects. A conventional method of UV-C exposure is the use of a collimated beam apparatus, which is assumed to employ sufficient uniform mixing while samples undergo UV-C exposure doses. In UV-C dose-response data it is assumed that the overall fluid is subjected to uniform dose distribution, while vigorous mixing takes place to provide accurate calculations (Akwu, et al., 2022). This assumption may not be applicable in some instances where the test fluid and micro-organisms utilized are different than described above. UV-C irradiation effectively inactivated *L. monocytogenes* ATCC 19115 in apple juice as may be seen in Fig. 5. The populations of *L. monocytogenes* were reduced by 1.20, 2.26, 3.48 log_10_ respectively at a UV-C dose level of 4, 8, 12 mJ·cm^-2^. The D_10_ value obtained overall in the apple juice was 3.53 mJ/cm^2^. The graph demonstrates that when the UV dosage is increased, the log reduction is increasing. Based on the rate constant (cm^2^/mJ) and D_10_ value of *L. monocytogenes*, 17.7 mJ.cm^-2^ dosage will be required to achieve 5 log_10_ reduction. Previously published literature data suggested that *L. monocytogenes* UV sensitivity lies between 3.24 mJ.cm^-2^ (Akwu, et al., 2022; Gunter-Ward, et al., 2017), 4 mJ.cm^-2^ log reduction (Akwu, et al., 2022; Lu, et al., 2010), and greater than 5 mJ.cm^-2^ (Akwu, et al., 2022; Matak, et al., 2005). D values that were higher than normal are possibly from lack of uniform dose delivery, which can be from poor mixing, causing huge UV gradients and inaccurate data. The FDA set the regulated UV-C dose at 40 mJ·.cm^-2^;which is predicted that inactivation treatments should result in a 5-log_10_ decrease of *S*. Muenchen, and *L. monocytogenes* in apple juice. To conclude, this system almost allowed accurate estimation of the delivered doses required to inactivate 5 log_10_ reductions of these pathogens taking into account the UV-C irradiance, absorption coefficient (1/cm) (Akwu, et al., 2022). Kinetic data shows that both pathogens are very sensitive to UV light at 279 nm wavelength. Overall, in this study, 20 mJ/cm^-2^ of exposure treatment is the optimized dosage for safer and higher quality product as also demonstrated in (Akwu, et al., 2022).

**Figure 4:**
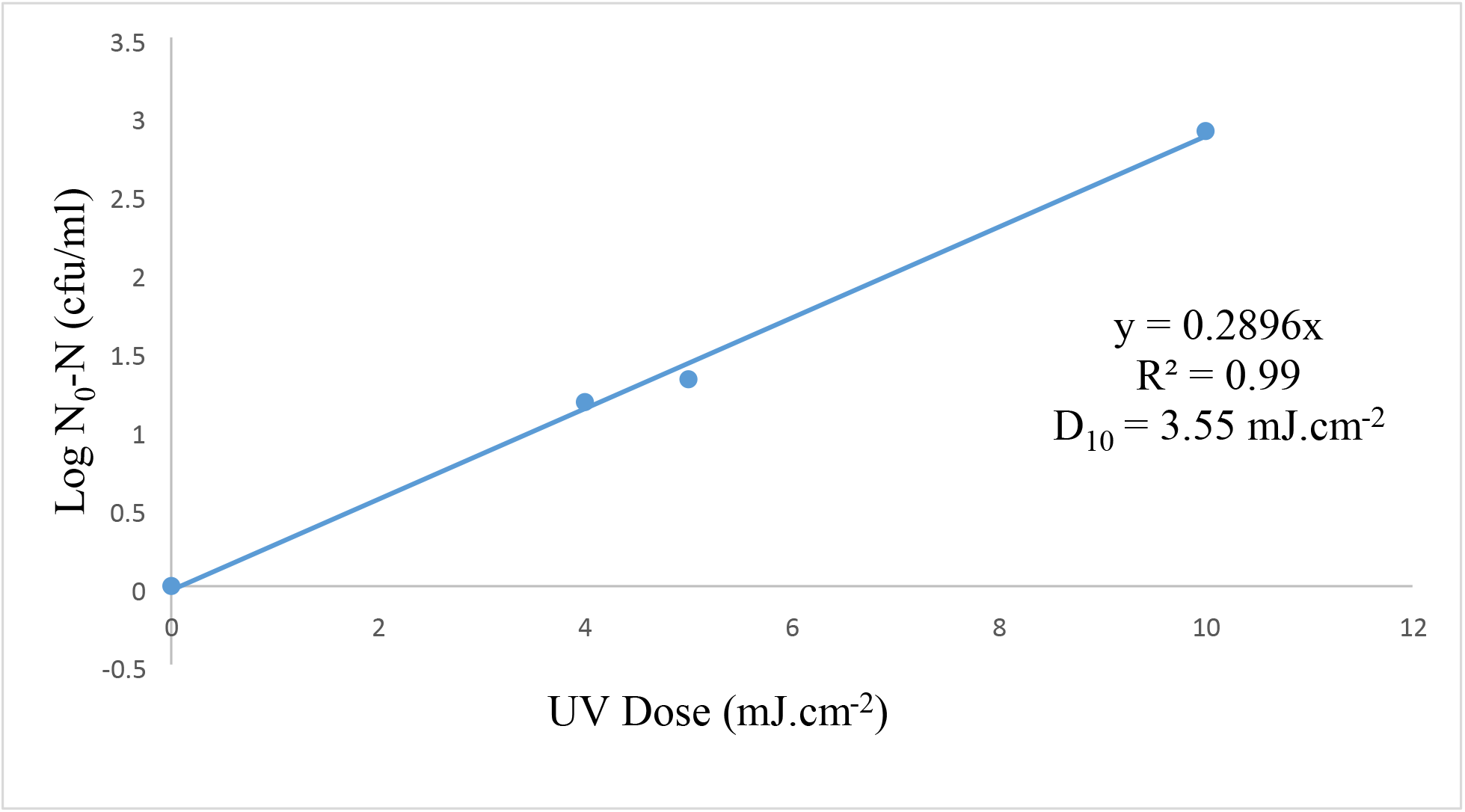
UV-C Inactivation of *Salmonella Muenchen* in apple juice using a collimated Light Emitting Diode UV system at 279 nm wave-length; the fluence intensity gradients were adjusted (i.e. optical properties of apple juice). Triplicate irradiations were performed for each dose; all replicates shown on plot, and values shown are averages of duplicate plating of each irradiated sample. Error bars represent range of data

**Figure 5:**
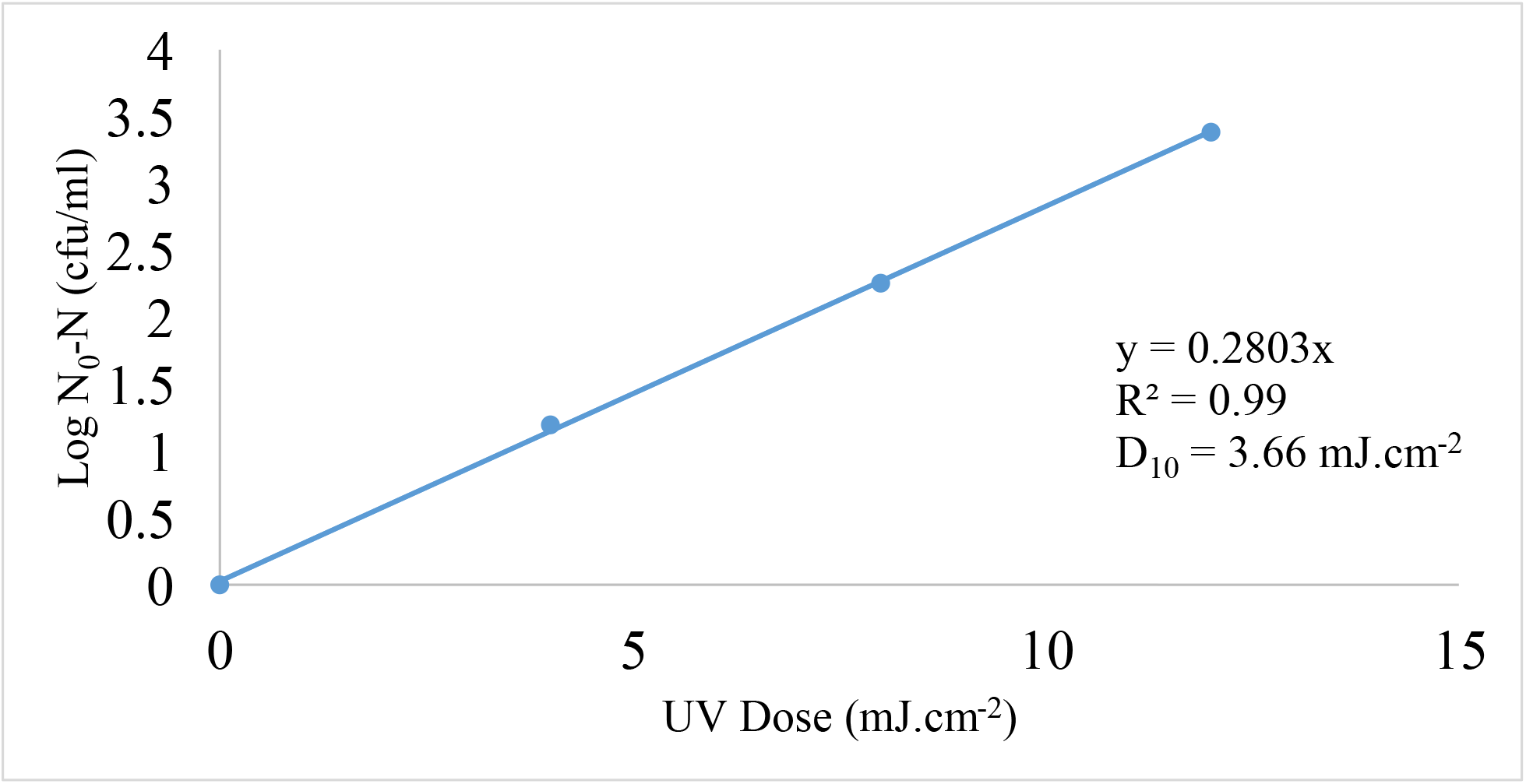
UV-C Inactivation of and *Listeria monocytogenes* in apple juice using a collimated Light Emitting Diode UV system at 279 nm wave-length; the fluence intensity gradients were adjusted (i.e. optical properties of apple juice). Triplicate irradiations were performed for each dose; all replicates shown on plot, and values shown are averages of duplicate plating of each irradiated sample. Error bars represent range of data

Figures 7 and 8 illustrates the effect of UV-C irradiation on the content of polyphenolic and vitamin compounds present in AJ. Figure 6 demonstrates chromatograms of chlorogenic acid and epicatechin respectively; chlorogenic acid has notably demonstrated as being the most abundant of polyphenolic compounds present in apple juice (Akwu, et al., 2022; Bender & Atalay, 2021; Islam, et al., 2016a; Islam, et al., 2016b; Eisele & Drake, 2005). Another abundant polyphenol in apple juice is epicatechin (Akwu, et al., 2022; Marcotte, et al., 2022; Bae, et al., 2020; Boyer & Liu, 2004). Concentration of epicatechin was observed to diminish as a function of UV dosage (p < 0.05) (Akwu, et al., 2022). At the maximum dosage level (160 mJ.cm^-2^), epicatechin and chlorogenic acid were significantly reduced by 98% and 76% respectively, similarly demonstrated in (Akwu, et al., 2022). Results suggested that epicatechin was relatively sensitive to UV-C dosage as compared to chlorogenic; as similarly found in Akwu et al., (2022). Overall, the data demonstrates an approximate 30% reduction at 20 mJ.cm^-2^ for both polyphenols (Akwu et al., 2022).

**Figure 6.**
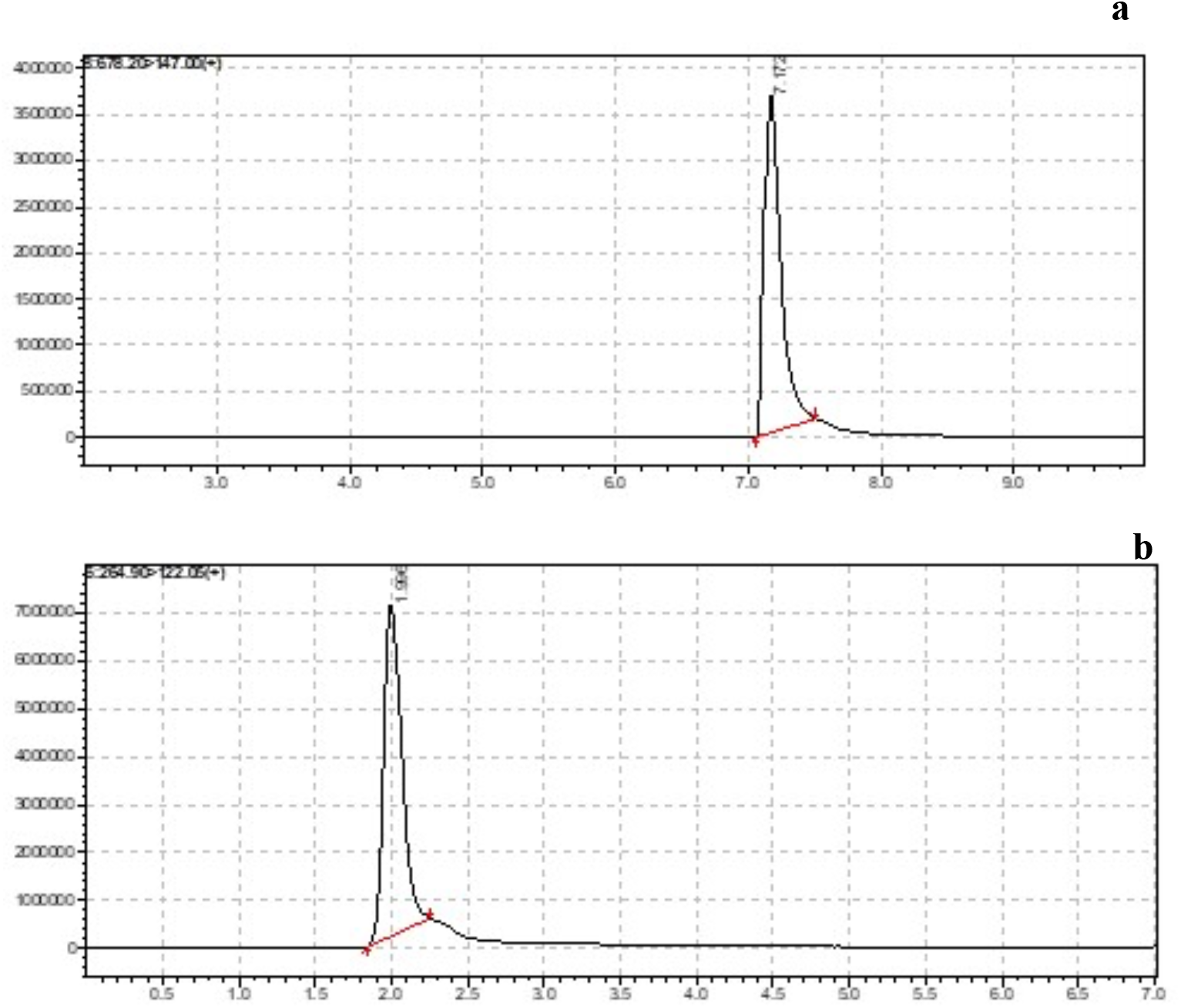
Representative Chromatograms of Vitamin B12 (a) and Thiamine (Vitamin B1) (b).

**Figure 7.**
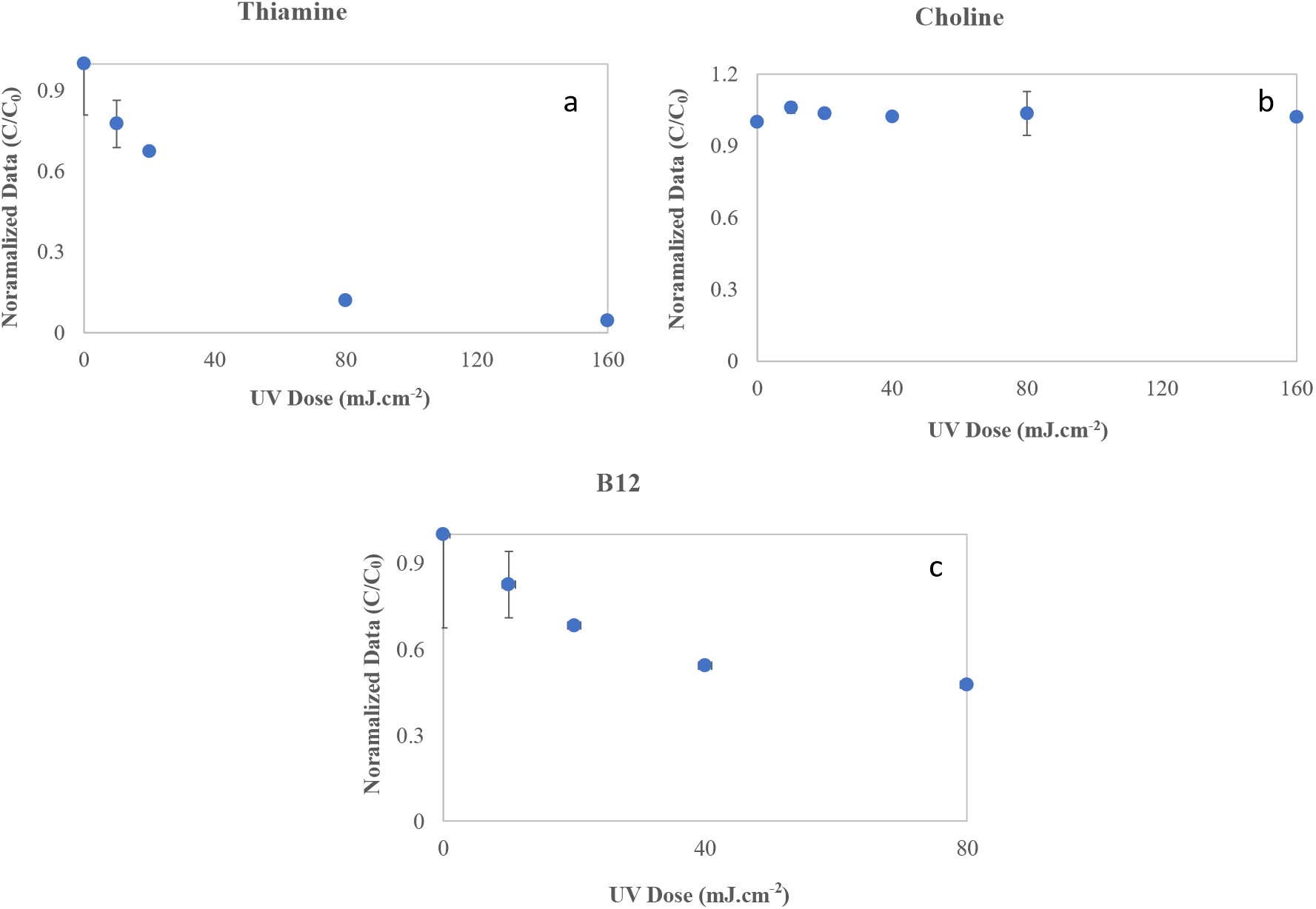
Effect of UV irradiation on the stability of vitamins [a. B12; b. Choline; c. Thiamine]. as a function of UV dose. The plots contain aggregated data from multiple experiments in which exposures were performed three times at each level. Error bars represent range of data

**Figure 8.**
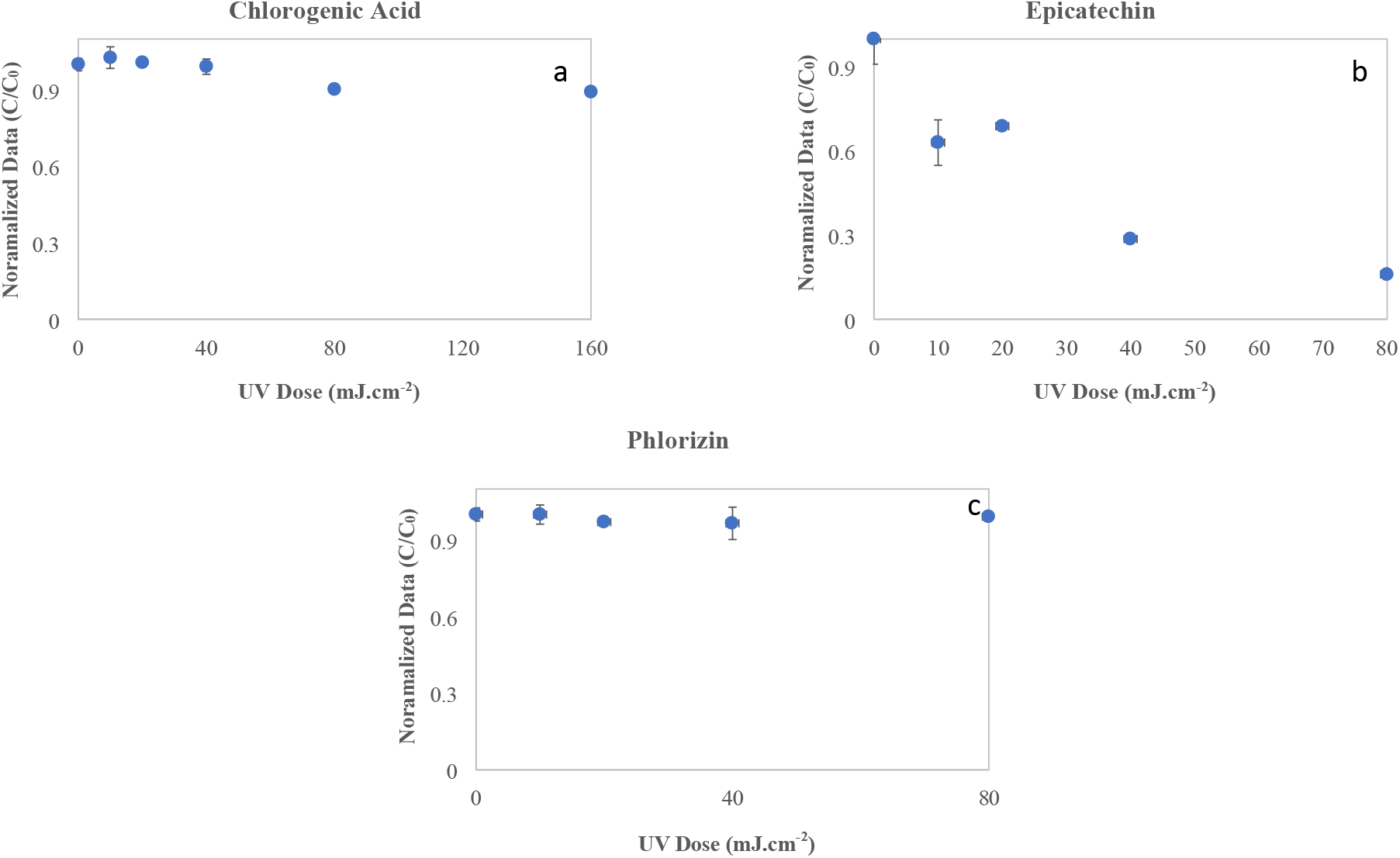
Effect of UV irradiation on the stability of polyphenols [a. Epicatechin; b. Chlorogenic acid; c. Phlorizin] as a function of UV dose. The plots contain aggregated data from multiple experiments in which exposures were performed three times at each level. Error bars represent range of data

Three vitamins and three polyphenols were identified and quantified in apple juice by LCMS/MS; all of the vitamins identified were significantly reduced (p < 0.05) as UV dosage was increased. Chlorogenic Acid, Epicatechin, Phlorizin, Thiamine (B1), Choline (B7), and B12. Thiamine (Stawny, et al., 2020; Whitfield, et al., 2018), and B12 (Juzeniene & Nizauskaite, 2013) are both photosensitive vitamins and therefore can demonstrate behaviors as photosensitizers; which is mainly due to double bond presence within their chemical structures. Additionally, this behavior can speed up other B vitamins oxidation processes (De Arrivetti et al., 2013). In this study, epicatechin concentrations in the samples decreased by 37%, 31%, 71%, and 84% at UV doses at 10, 20, 40, and 80 mJ.cm^-2^, respectively. Epicatechin was rapidly degraded by UV-C exposures. Similar findings were reported in three other studies (Akwu et al., 2022; (Islam, et al., 2016a; Islam, et al., 2016b; Akwu et al., 2022). Based on the observed concentration reductions, it demonstrated that all vitamins subjected to the UV-C exposure demonstrated sensitivity in apple juice and as a result they decreased significantly. The mechanism responsible for vitamin degradation during UV irradiation is possibly from a couple of different chemical-based reactions and mechanisms to include photo-oxidation, thermal degradation, (Islam, et al., 2016b; Sheraz et al., 2014; Akwu et al., 2022). In contrast, chlorogenic acid retained very well in the UV exposed samples, and the thiamine was significantly reduced following exposure to UV-C. Our study demonstrated that UV-C irradiation caused significant changes in vitamins and polyphenols, and it is quite indicative that UV irradiation is capable of changing a number of chemical food constituents similarly demonstrated in Akwu et al., (2022). To the best of the authors’ knowledge, this is the first paper that demonstrates the efficacy of UVC 279 nm inactivating *Salmonella* Muenchen, and *Listeria monocytogenes* in highly turbid apple juice and retaining the quality between 20 - 40 mJ.cm^-2^ dosage.

## Conclusion

UV-C irradiation was successfully applied to inactivate the microbial populations in apple juice using a near collimated beam device operating at 279 nm wavelength. This study found that UVC irradiation treatment at low doses (≈25 mJ·cm^-2^) could be used to achieve 5-log_10_ inactivation of *Salmonella* Muenchen, and *Listeria monocytogenes* strains. In addition, the D_10_ values of 4.12 and 3.56 mJ·cm^-2^ were obtained for *Salmonella enterica* serovar Muenchen ATCC BAA 1764, and *Listeria monocytogenes* ATCC 19115. The inactivation kinetics of these tested microorganisms were best described by log linear kinetics. UV-C irradiation induced minor reductions in the concentration of vitamins and polyphenols in apple juice at the FDArecommended fluency of 40 mJ cm^-2^ of pasteurized equivalent dose. Chlorogenic acid was reduced to 56%, at 80 mJ/cm2 whereas 12% reduction was observed at 40 mJ/cm^2^. The decrease in the concentration of antioxidants is dependent on various intrinsic factors such as pH, presence of light sensitive aromatic chemical compounds and formation of non-specific reactive oxygen species Epicatechin was observed to be relatively stable as a function of UV dosage. Scale-up of the UV-C LED device (flow-through system), spore inactivation studies, and sensory evaluation of UV-C treated apple juice will be subject of further investigations. The lab is developing a scaleup model and its efficacy in inactivating microorganisms and other spores in apple juice on a larger scale will be subject to future investigation.

## Acknowledgments

The authors would like to thank Mrs. Yvonne Myles, Dr. Jerwzy Mierzwa, Mr. Vybhav Gopisetty, and Ms. Judy Stanley for providing valuable guidance in this project.

## References

Administration, U.S.F.D. (2020). United States Food and Drug Administration. Ultraviolet (UV) Radiation. https://www.fda.gov/radiation-emitting-products/tanning/ultraviolet-uv-radiation.

Akwu, A.S., A. Patras, B. Pendyala, A. Kurup, F-C. Chen, and M.J. Vergne. (2022). Effect of germicidal short wave-length ultraviolet light on the polyphenols, vitamins, and microbial inactivation in highly opaque apple juice. bioRxiv, 87, 123–456. https://doi.org/10.1101/2022.07.29.502038

Aprea, E., M. Charles, I. Endrizzi, M.L. Corollaro, E. Betta, F. Biasioli, and F. Gasperi. (2017). Sweet taste in apple: the role of sorbitol, individual sugars, organic sugars and volatile compounds. Scientific Reports, 7, 44950. https://doi.org/10.1038/srep44950

Assatarakul, K.; Churey, J. J.; Manns, D. C.; & Worobo, R. W. (2012). Patulin reduction in apple juice from concentrate by UV radiation and comparison of kinetic degradation models between apple juice and apple cider. Journal of Food Protection, 75, 717–724. https://doi.org/10.4315/0362-028X.JFP-11-429

Bae, J., Kim, N., Shin, Y., Kim, S.-Y., & Kim, Y.-J. (2020). Activity of catechins and their applications. Biomedical Dermatology, 4 (8). https://doi.org/10.1186/s41702-020-0057-8

Beck, S.E., Ryu, H., Boczek, L.A., Cashdollar. J.L., Jeanis, K.M., Rosenblum, J.S., Lawal, O.R. & Linden, K.G. (2017). Evaluating UV-C LED disinfection performance and investigating potential dual-wavelength synergy. Water Resource, 109, 207–216. https://doi.org/10.1016/j.watres.2016.11.024

Bender, O., & Atalay, A. (2021). Chapter 28-Polyphenol chlorogenic acid, antioxidant profile, and breast cancer. Cancer (Second Edition): Oxidative Stress and Dietary Antioxidants. Academic Press, pp 311–321. https://doi.org/10.1016/B978-0-12-819547-5.00028-6

Bhullar, M.S., A. Patras, A. Kilonzo-Nthenge, B. Pokharel, and M. Sasges. (2019). Ultraviolet inactivation of bacteria and model viruses in coconut water using a collimated beam system. Food Science and Technology International, 25,7, 562–572. https://doi.org/10.1177/1082013219843395

Bolton, J.R. & Linden, K.G. (2003). Standardization of Methods for Fluence (UV Dose) Determination in Bench-Scale UV Experiments. Journal of Environmental Engineering, 129 (3), 209–216. https://doi.org/10.1061/ASCE0733-93722003129:3209

Boyer, J.; & R.H. Liu. 2004. Apple phytochemicals and their health benefits. Nutr. J. 2004, 3, 1–15. https://doi.org/10.1186/1475-2891-3-5

Caminiti, I. M.; Palgan, I.; Muňoz, A.; Noci, F.; Whyte, P.; Morgan, D. J.; Cronin, D. A.; Lyng, J. G. The effect of ultraviolet light on microbial inactivation and quality attributes of apple juice. Food Bioprocess Technol. 2012, 5, 680–686. https://doi.org/10.1007/s11947-010-0365-x.

Center for Food Safety and Applied Nutrition (C. F. S. A. N.), United States Food and Drug Administration, United States Department of Health and Human Services. (2000, March 29). Kinetics of Microbial Inactivation for Alternative Food Processing Technologies: IFT/FDA Contract No. 223-98-2333 [PDF file]. Retrieved from https://www.fda.gov/files/food/published/Kinetics-of-Microbial-Inactivation-for-AlternativeFood-Processing-Technologies.pdf.

Chandra, S., A. Patras, B. Pokharel, R.R. Bansode, A. Begum, and M. Sasges. 2017. Patulin degradation and cytotoxicity evaluation of UV irradiated apple juice using human peripheral blood mononuclear cells. Journal of Food Process Engineering, 40, 6, e12586. https://doi.org/10.1111/jfpe.12586.

Chevremont, A.-C., Farnet, A.-M., Coulomb, B., & Boudenne, J.-L.. 2012. Effect of coupled UV-A and UV-C LEDs on both microbiological and chemical pollution of urban wastewaters. Science of the Total Environment, 426, 304–310. https://doi.org/10.1016/j.scitotenv.2012.03.043

Cousin, F.J., R. Le Guellec, M. Schlusselhuber, M. Dalmasso, J.-M. Laplace, and M. Cretenet. 2017. Microorganisms in Fermented Apple Beverages: Current Knowledge and Future Directions. Microorganisms, 5,3,39. https://doi.org/10.3390/microorganisms5030039

Dai, T. Vrahas, M.S., Murray, C.K., & Hamblin, M.R. (2012). Ultraviolet C irradiation: an alternative antimicrobial approach to localized infections? Expert Review of Anti-infective Therapy, 10(2), 185–195. https://doi.org/10.1586/eri.11.166

De Arrivetti, O. R., L.; Scurachio, R. S.; Santos, W. G.; Papa, T. B.R.; Skibsted, L. H.; Cardoso, D. R. Photooxidation of Other B Vitamins as Sensitized by Riboflavin. J. Agric. Food Chem. 2013, 61,7615–7620. https://doi.org/10.1021/jf402123d

Delorme, M.M., Guimarã, J.T., Coutinho, N.M., Balthazar, C.F., Rocha, R.S., Silva, R., Margalho, L.P., Pimentel, T.C., Silva, M.C., Freitas, M.Q., Granato, D., Sant’Ana, A.S., Duart, M.C.K.H., Cruz, A.G. 2020. Ultraviolet radiation: An interesting technology to preserve quality and safety of milk and dairy foods. Trends in Food Science & Technology, 102, 146–154. https://doi.org/10.1016/j.tifs.2020.06.001

Dimitrovski, D., E. Velickova, T. Langerholc, and E. Winkelhausen. 2015. Apple juice as a medium for fermentation by the probiotic *Lactobacillus plantarum* PCS 26 strain. Annals of Microbiology, 65, 2161–2170. https://doi.org/10.1007/s13213-015-1056-7

Du, G., Y. Zhu, X. Wang, J. Zhang, C. Tian, L. Liu, Y. Meng, and Y. Guo. 2019. Phenolic composition of apple products and by-products based on cold pressing technology. Journal of Food Science and Technology, 56,3,1389–1397. https://doi.org/10.1007/s13197-019-03614-y

Eisele, T.A. and S.R. Drake. 2005. The partial compositional characteristics of apple juice from 175 apple varieties. J. Food Compos. Anal. 18, 213–221. https://doi.org/10.1016/j.jfca.2004.01.002

Gehr, R. 2007. Collimated beam tests: their limitations for assessing wastewater disinfectability by UV, and a proposal for an additional evaluation parameter. Journal of Environmental Engineering and Science, 6 (3). https://doi.org/10.1139/s06-032

Gopisetty, V.V.S., Patras, A., Pendyala, B., Kilonzo-Nthenge, A., Ravi, R., Pokharel, B., Zhang, L., Si, H., & Sasges, M. 2019. UV-C irradiation as an alternative treatment technique: Study of its effect on microbial inactivation, cytotoxicity, and sensory properties in cranberry-flavored water. Innovative Food Science & Emerging Technologies, 52, 66–74. https://doi.org/10.1016/j.ifset.2018.11.002

Gunter-Ward, D., A. Patras, M.S. Bhullar, A. Kilonzo-Nthenge, B. Pokharel, and M.R. Sasges. 2017. Efficacy of ultraviolet (UV-C) light in reducing foodborne pathogens and model viruses in skim milk. Journal of Food Processing and Preservation, 42,1. http://doi.org/10.1111/jfpp.13485.

Heckman, C.J., Chandler, R., Kloss, J.D., Benson, A., Rooney, D., Munshi, T., Darlow, S.D., Perlis, C., Manne, S.L. & Oslin, D.W. 2013. Minimal Erythema Dose (MED) Testing. Journal of Visualized Experiments, 75, 50175. https://doi.org/10.3791/50175.

Heinmaa, L., U. Moor, P. Põldma, P. Raudsepp, U. Kidmose, and R.L. Scalzo. 2016. Content of health-beneficial compounds and sensory properties of organic apple juice as affected by processing technology. LWT Food Sci Technol., 85,372–379. https://doi.org/10.1016/j.lwt.2016.11.044

Islam, M. S.; Patras, A.; Pokharel, B.; Vergne, M.; Sasges, M.; Begum, A.; Rakariyatham, K.; Pan, C.; and Xiao, H. Effect of UV Irradiation on the Nutritional Quality and Cytotoxicity of Apple Juice. J. Agric. Food Chem. 2016b, 64, 41, 7812–7822 https://doi.org/10.1021/acs.jafc.6b02491

Islam, M. S.; Patras, A.; Pokharel, B.; Wu, Y.; Vergne, M. J.; Shade, L.; Xiao, H.; Sasges, M. UV-C irradiation as an alternative disinfection technique: Study of its effect on polyphenols and antioxidant activity of apple juice. Innovative Food Sci. Emerging Technol. 2016a, 34, 344–351 https://doi.org/10.1016/j.ifset.2016.02.009

Jimenez-Garcia et al., S.N., M. A. Vazquez-Cruz, L. Garcia-Mier, L.M. Contreras-Medina, R.G. Guevara-González, J. F. Garcia-Trejo, and A.A. Feregrino-Perez. 2018. Phytochemical and Pharmacological Properties of Secondary Metabolites in Berries. Therapeutic Foods: Handbook of Food Bioengineering, Chapter 13, 397–427. https://doi.org/10.1016/B978-0-12-811517-6.00013-1.

Juzeniene, A., & Nizauskaite, Z. 2013. Photodegradation of cobalamins in aqueous solitions and in human blood. Journal of Photochemistry and Photobiology, B, Biology. 122, 7–14. https://doi.org/10.1016/j.jphotobiol.2013.03.001.

Karasawa, M.M.G., and C. Mohan. 2018. Fruits as Prospective Reserves of bioactive Compounds: A Review. Nat. Prod.Bioprospect, 8,5, 335–346. https://doi.org/10.1007/s13659-018-0186-6.

Kebbi, Y., Muhammad, A.I., Sant’Ana, A.S., do Prado-Silva, L., Liu, D., & Ding, T. 2020. Recent advances on the application of UV-LED technology for microbial inactivation: Progress and mechanism. Comprehensive Reviews in Food Science and Food Safety, 19(6), 3501–3527. https://doi.org/10.1111/1541-4337.12645.

Kidoń, M. and J. Grabowska. 2021. Bioactive compounds, antioxidant activity, and sensory qualities of red-fleshed apples dried by different methods. LWT Food Science and Technology, 136,2, 110302. https://doi.org/10.1016/j.lwt.2020.110302.

Kim, S.-J., Kim, D.-K., & Kang, D.-H. 2015. Using UVC Light-Emitting Diodes at Wavelengths of 266 to 279 Nanometers to Inactivate Foodborne Pathogens and Pasteurize Sliced Cheese. Applied Environmental Microbiology, 82(1), 11–17. https://doi.org/10.1128/AEM.02092-15

Koutchma, T. 2019. Ultraviolet Light in Food Technology: Principles and Applications. 2^nd^ Edition, Boca Raton, CRC Press, https://doi.org/10.1201/9780429244414

Kuo, J., Asce, M., Chen, C.-L., & Nellor, M. 2003. Standardized collimated beam testing protocol for water/wastewater ultraviolet disinfection. Journal of Environmental Engineering, 129, 774–779. https://doi/org/10.1061/(ASCE)0733-9372(2003)129:8(774)

Kurup, A.H., A. Patras, B. Pendyala, M.J. Vergne, R.R. Bansode. 2022. Evaluation of Ultraviolet-Light (UV-A) Emitting Diodes Technology on the Reduction of Spiked Aflatoxin B_1_ and Aflatoxin M_1_ in Whole Milk. Food and Biopress Technology, 15, 165–176 https://doi.org/10.1007/s11947-021-02731-x

Kusuma, P., Pattison, P.M., & Bugbee, B. 2020. From physics to fixtures to food: current and potential LED efficacy. Horticulture Research, 7,56. https://doi.org/10.1038/s41438-020-0283-7

Liu, R.H. 2004. Potential synergy of phytochemicals in cancer prevention. Mechanism of action. The Journal of Nutrition, 134, 3479S–34855S. https://doi.org/10.1093/jn/134.12.3479S.

Lu, G., Li, C., Liu, P., Cui, H., Yao, Y., & Zhang, Q. (2010). UV inactivation of microorganisms in beer by novel thin-film apparatus. Food Control, 21(10), 1312–1317. https://doi.org/10.1016/j.foodcont.2010.03.007

Marcotte, B.V., M. Verheyde, S. Pomerleau, A. Doyen, and C. Couillard. 2022. Health Benefits of Apple Juice Consumption: A Review of Interventional Trials on Humans. Nutrients, 14(4): 821. https://doi.org/10.3390/nu14040821

Matak, K. E., Churey, J. J., Worobo, R. W., Sumner, S. S., Hovingh, E., Hackney, C. R., & Pierson, M. D. (2005). Efficacy of UV-C light for the reduction of Listeria monocytogenes in goat’s milk. Journal of Food Protection, 68, 2212–2216. https://doi/org/10.4315/0362-028x-68.10.2212.

McSharry, S., Koolman, L., Whyte, P., & Bolton, D. (2022). Inactivation of *Listeria monocytogenes* and *Salmonella Typhimurium* in beef broth and on diced beef using an ultraviolet light emitting diode (UV-LED) system. Lebensmittel-Wissenschaft & Technologie (LWT), Food Science and Technology, 158, 113150. https://doi.org/10.1016/j.lwt.2022.113150

Medicine, J.H. (2022). Johns Hopkins Medicine, Health: Ultraviolet Radiation. https://www.hopkinsmedicine.org/health/conditions-and-diseases/ultraviolet-radiation.

Miller, D. N., Bryant, J. E., Madsen, E. L., & Ghiorse, W. C. (1999). Evaluation and optimization of DNA extraction and purification procedures for soil and sediment samples. Applied and Environmental Microbiology, 65(11), 4715–4724. https://doi.org/10.1128/AEM.65.11.4715-4724.1999.

Muñoz, A., I.M. Caminiti, I. Palgan, G. Pataro, F. Noci, D.J. Morgan, D.A. Cronin, P. Whyte, G. Ferrari, and J.G. Lyng. 2012. Effects on *Escherichia coli* inactivation and quality attributes in apple juice treated by combinations of pulsed light and thermosonication. Food Research International, 45,1, 299–305. https://doi.org/10.1016/j.foodres.2011.08.020

Nyhan, L., Przyjalgowski, M., Lewis, L., Begley, M., & Callanan, M. 2021. Investigating the Use of Ultraviolet Light Emitting Diodes (UV-LEDs) for the Inactivation of Bacteria in Powdered Food Ingredients. Foods, 10(4), 797. https://doi.org/10.3390/foods10040797

Organization, W.H. (2016). World Health Organization. Radiation: Ultraviolet (UV) radiation. https://www.who.int/news-room/questions-and-answers/item/radiation-ultraviolet-(uv).

Patras, A., B.K. Tiwari, and N.P. Brunton. 2011. Influence of blanching and low temperature preservation strategies on antioxidant activity and phytochemical content of carrots, green beans and broccoli. LWT - Food Science and Technology, 299–306 https://doi.org/10.1016/j.lwt.2010.06.019

Patras, A., Ricketts, J., Pendyala, B., & Godwin, S. (2020). UV-C Light Ensuring Safety and Quality of Beverages. 1 – 2. https://www.tnstate.edu/extension/documents/UV%20Light%20Fact%20Sheet%20FSBE-01-2020-1.pdf. (2020, May 18). UV-C Light Ensuring Safety and Quality of Beverages [PDF file].

Pendyala, B., Patras, A. Gopisetty, V.V.S., & Sasges, M. 2021. UV-C inactivation of microorganisms in a highly opaque model fluid using a pilot scale ultra-thin film annular reactor: Validation of delivered dose. Journal of Food Engineering, 294(14),110403. https://doi.org/10.1016/j.jfoodeng.2020.110403

Pina-Pérez, M.C., D. Rodrigo, and A. Martinez. 2015. Using natural antimicrobials to enhance the safety and quality of fruit-and vegetable-based beverages. Handbook of Natural Antimicrobials for Food Safety and Quality, Chapter 16, 347–363. https://doi.org/10.1016/B978-1-78242-034-7.00016-5

Prasad, A., Du, L., Zubair, M., Subedi,S., Ullah, A., & Roopesh, M.S. 2020. Applications of Light Emitting Diodes (LEDs) in Food Processing and Water Treatment. Food Engineering Reviews, 12, 268–289. https://doi.org/10.1007/s12393-020-09221-4

Prevention, C.D.C. (2022). Centers for Disease Control and Prevention: National Center for Environmental Health. UV Radiation. https://www.cdc.gov/nceh/features/uv-radiationsafety/index.html#:~:text=Ultraviolet%20(UV)%20radiation%20is%20a,The%20sun

Qualls, R. G., Flynn, M. P., & Johnson, J. D. 1983. The Role of Suspended Particles in Ultraviolet Disinfection. Water Pollution Control, 55, (10),1280–1285. https://doi.org/10.2307/25042084

Rahman, M.S. 2020. UV Light in Food Preservation (Handbook of Food Preservation),3^rd^ Edition, CRC Print.

Rasooly, R., Magoz, Z., Luo, J., Do, P., Hernlem, B. 2018. Sensitive low-cost CCD-based detector for determination of UV-LED water microbial disinfection. Desalination and Water Treatment, 118, 120–125. https://doi.org/10.5004/dwt.2018.22667

Ribeiro, J.A. E. dos Santos Pereira, C. de Oliveira Raphaelli, M. Radünz, T.M. Camargo, F.I.G. da Rocha Concenço, R.F.F. Cantillano, Â.M. Fiorentini, and L. Nora. 2021. Application of prebiotics in apple products and potential health benefits. Journal of Food Science and Technology, 59, 1249–1262. https://doi.org/10.1007/s13197-021-05062-z

Sauceda-Gálvez, J.N., M. Martinez-Garcia, M.M. Hernández-Herrero, R. Gervilla, and A.X Roig-Sagués. 2021. Short Wave Ultraviolet Light (UV-C) Effectiveness in the Inactivation of Bacterial Spores Inoculated in Turbid Suspensions and in Cloudy Apple Juice. Beverages, 7, 11. https://doi.org/10.3390/beverages7010011.

Sheraz, M.A., Kazi, S.H., Ahmed, S., Anwar, Z., & Ahmad, I. (2014). Photo, thermal, and chemical degradation of riboflavin. Beilstein Journal of Organic Chemistry 10, 1999–2012. https://doi.org/10.3762/bjoc.10.208

Smeriglio, A., D. Barreca, E. Bellocco, D. Trombetta. 2017. Proanthocyanidins and hydrolysable tannins: occurrence, dietary intake and pharmacological effects. British Journal of Pharmacology, 174, 11, 1244–1262. https://doi.org/10.1111/bph.13630

Sommer, R., Lhotsky, M., Haider, T., & Cabaj, A. 2000. UV inactivation, liquid-holding recovery, and photoreactivation of Escherichia coli O157 and other pathogenic Escherichia coli strains in water, Journal of Food Protection, 63 (8), 1015–1020. https://doi.org/10.4315/0362028x-63.8.1015.

Stanley, J., A. Patras, B. Pendyala, M.J. Vergne, & R.R. Bansode. 2020. Performance of a UV-A LED system for degradation of aflatoxins B_1_ and M_1_ in pure water: kinetics and cytotoxicity study. Scientific Reports, 10, 1, 1–12. https://doi.org/10.1038/s41598-020-70370-x

Stawny, M., Gostyńska, A., Olijarczyk, R., Jelińska, A., & Ogrodowczyk, M. 2020. Stability of high-dose thiamine in parenteral nutrition for treatment of patients with Wernicke’s encephalopathy, 39 (9), 2929–2932. Clinical Nutrition (Edinburg, Scotland). https://doi.org/10.1016/j.clnu.2019.12.003

Sulaiman, A., M. Farid, and F.V.M. Silva. 2016. Quality stability and sensory attributes of apple juice processed by thermosonication, pulsed electric field and thermal processing. Food Science and Technology International, 23,3, 265–276. https://doi.org/10.1177/1082013216685484

Swamy, G.J., K. Muthukumarappan, and S. Asokapandian. 2018. Ultrasound for Fruit Juice Preservation. Fruit Juices Extraction, Composition, Quality and Analysis. Chapter 23, 451–462. https://doi.org/10.1016/B978-0-12-802230-6.00023-0

Teleszko, M. & Wojdyło, A. 2015. Comparison of phenolic compounds and antioxidant potential between selected edible fruits and their leaves. J. Funct. Foods 14, 736–746. https://doi.org/10.1016/j.jff.2015.02.041.

Torkamani, A.E. & Niakousari, M. 2011. Impact of UV-C light on orange juice quality and shelf life. International Food Research Journal, 18,4, 1265.

Tosa, K., & Hirata, T. (1999). Photoreactivation of enterohemorrhagic Escherichia coli following UV disinfection. Water Research, 33(2), 361–366. https://doi.org/10.1016/S0043-1354(98)00226-7

Unluturk, S.; Atilgan, M. R.; Baysal, A. H.; Unluturk, M. Modeling inactivation kinetics of liquid egg white exposed to UV-C irradiation. Int. J. Food Microbiol. 2010, 142, 341–347. https://doi.org/10.1016/j.ijfoodmicro.2010.07.013.

Usaga, J., D.C. Manns, C.I. Moraru, R.W. Worobo, O. Padilla-Zakour. 2017. Ascorbic acid and selected preservatives influence effectiveness of UV treatment of apple juice. LebensmittelWissenschaft und-technologie, 75. https://doi.org/10.1016/j.lwt.2016.08.037.

Van der Sluis, A.A.; Dekker, M.; Skrede, G.; & Jongen, W.M.F. Activity and concentration of polyphenolic antioxidants in apple juice effect of existing methods. J. Agric. Food Chem. 2002, 50, 7211–7219. https://doi.org/10.1021/jf020115h

Vashisht, P., B. Pendyala, A. Patras, V.V.S. Gopisetty, R. Ravi. 2022. Pilot scale study on UV-C inactivation of bacterial endospores and virus particles in whole milk: evaluation of system efficiency and product quality. bioRxiv, 1–35. https://doi.org/10.1101/2022.01.07.475436.

Whitfield, K.C., M.W. Bourassa, B. Adamolekun, G. Bergeron, L. Bettendorff, K.H. Brown, L. Cox, A. Fattal-Valevski, P.R. Fischer, E.L. Frank, L. Hiffler, L.M. Hlaing, M.E. Jefferds, H. Kapner, S. Kounnavong, M.P.S. Mousavi, D.E. Roth, M-N. Tsaloglou, F. Wieringa, and G.F. Combs,Jr. 2018. Thiamine deficiency disorders: diagnosis, prevalence, and a roadmap for global control programs. Annals of the New York Academy of Sciences, 1430,1, 3–43. https://doi.org/10.1111/nyas.13919

Wilson, B. R., Roessler, P.F., van Dellen, E., Abbaszadegan, M., & Gerba, C.P. (1992). Coliphage MS2 as a UV water disinfection efficacy test surrogate for bacterial and viral pathogens. Proceedings of the American Water Works Association Water Quality Technology Conference, Toronto, Canada.

Włodarska, K., K. Pawlak-Lemańska, T. Górecki, and E. Sikorska. 2019. Factors Influencing Consumers’ Perceptions of Food: A Study of Apple Juice Using Sensory and Visual Attention Methods. Foods, 8, 11, 545. https://doi.org/10.3390/foods8110545

Wojdyło, A., P. Nowicka, I.P. Turkiewicz, K. Tkacz, and F. Hernandez. 2021. Comparison of bioactive compounds and health promoting properties of fruits and leaves of apple, pear, and quince. Scientific Reports, 11, 20253. https://doi.org/10.1038/s41598-021-99293-x.

Worobo, R. 1999. Efficacy of the CiderSure 3500 Ultraviolet light unit in apple cider. CFSAN Apple cider food safety control workshop (Washington D.C., USA).

Wu, D, You, H, Zhang, R, Chen, C., & Lee, D-J. (2011) Ballast waters treatment using UV/Ag-TiO2+ O3 advanced oxidation process with *Escherichia coli* and *Vibrio alginolyticus* as indicator microorganisms. Chemical Engineering Journal 174(2-3): 714–718. https://doi.org/10.1016/j.cej.2011.09.087

Xiang, Q., Fan, L., Zhang, R., Ma,Y., Liu, S., & Bai, Y. (2020). Effect of UVC light-emitting diodes on apple juice: Inactivation of *Zygosaccharomyces rouxii* and determination of quality. Food Control, 107082. https://doi.org/10.1016/j.foodcont.2019.107082

Yaun, B.R., Sumner, S.S., Eifert, J.D., & Marcy, J.E. (2003). Response of Salmonella and Escherichia coli O157:H7 to UV Energy, Journal of Food Protection, 66(6), 1071–1073. https://doi/org/10.4315/0362-028x-66.6.1071

Yang, J.-H., Wu, U.-I., Tai, H.-M., & Sheng, W.-H. (2019). Effectiveness of an ultraviolet-C disinfection system for reduction of healthcare-associated pathogens. Journal of Microbiology, Immunology and Infection, 52(3), 487–493. https://doi.org/10.1016/j.jmii.2017.08.017

Yin, R., Dai, T., Avci, P., Jorge, A.E.S., de Melo, W.C.M.A., Vecchio, D., Huang, Y.-Y., Gupta, A., & Hamblin, M.R. (2013). Light based anti-infectives: ultraviolet C irradiation, photodynamic therapy, blue light, and beyond. Current Opinion in Pharmacology, 13(5). https://doi.org/10.1016/j.coph.2013.08.009

Zhang, S., C. Hu, Y. Guo, X. Wang, and Y. Meng. (2021). Polyphenols in fermented apple juice: Beneficial effects on human health. Journal of Functional Foods, 104294. https://doi.org/10.1016/j.jff.2020.104294

